# M1 protein expression determines niche-specific spread of *Streptococcus pyogenes*

**DOI:** 10.64898/2026.02.11.705257

**Authors:** Kristin K Huse, Elita Jauneikaite, Nicola N Lynskey, Lucy C Reeves, Mark Reglinski, Matthew K Siggins, Claire E Turner, Virginia Tajadura-Ortega, Yuan Chen, Antonio Di Maio, Wengang Chai, Yan Liu, Lucy E Lamb, Shiranee Sriskandan

## Abstract

Invasive infections with *Streptococcus pyogenes* account for significant mortality. Cell surface M protein is believed to be essential for *S. pyogenes* virulence, with critical roles in dissemination and immune evasion. In contrast, the role of secreted M protein in infection is less clear. Here, we assessed the role of M protein in streptococcal colonisation, persistence, and invasion in respiratory and soft tissue murine infection models, and report that while cell-surface M protein is required to establish nasopharyngeal infection, it does not drive invasive infection. Cell-surface M protein was important for binding of major host matrix proteins and growth in immune human blood, but soluble M protein was the predominant determinant of lymphatic survival, tissue spread, and systemic infection in mice. A mutant *emm*1 strain which only expresses secreted M protein exhibited enhanced survival in the lymph node niche and overall competitive advantage over the wildtype parent strain in co-infections. Absence of M protein at the bacterial cell surface led to enhanced hyaluronan binding, providing explanation for homing to the draining lymph node. Together, these findings highlight potential nuance in the role of M protein that may impact vaccine approaches targeting M protein.

**Author Summary:** *S. pyogenes* is a significant cause of illness, both mild and lethal, worldwide. Despite this, there are currently no approved vaccines. Many vaccine candidates are based on the M protein, a surface expressed anti-phagocytic factor that is thought to be critical for the ability of *S. pyogenes* to evade the host immune system. The unexpected selection of an M1 protein mutant strain following *in vivo* soft tissue infection raised many questions for us. If M protein is meant to be the critical factor in avoiding immune killing, how was this strain able to persist? One explanation could be that, although the strain was negative for M1 protein on the bacterial surface, it still secreted a truncated version of the protein with potential inflammatory effects. Importantly, the strain was selected in the draining lymph node pointing to a niche-specific advantage. We showed that cell surface-associated M1 protein, although required to establish nasopharyngeal infection, was not required in soft tissue infection, and that expression of a soluble, truncated form of M1 protein gave a survival advantage over the wild-type strain expressing cell wall-associated M1 protein. Furthermore, the absence of cell surface M protein led to enhanced hyaluronan binding providing explanation for lymphatic niche homing.

## Introduction

*Streptococcus pyogenes* (also known as group A *Streptococcus*) is a bacterial pathogen that is remarkable in its ability to cause a very wide range of disease manifestations (1). Despite a high burden of disease and mortality from both invasive infections and rheumatic heart disease [2], no licensed vaccine currently exists for *S. pyogenes*. The need for a vaccine has been recently underlined by the upsurge of invasive *S. pyogenes* disease in several countries following relaxation of COVID-19 related restrictions, alongside emergence of the new M1_UK_ lineage [3–6].

A crucial step in *S. pyogenes* establishing any type of infection is attachment to tissues, followed by evasion of the host immune response, against which *S. pyogenes* has a large arsenal of virulence factors. One of the key players in immune evasion is the surface-anchored M protein, encoded by the *emm* gene. A long history of research points to a vital role for M protein in *S. pyogenes* pathogenesis and as a potential vaccine target. M protein interacts with a wide array of host factors such as complement inhibitors, fibrinogen, fibronectin, plasminogen and immunoglobulin [7]. The binding of fibrinogen by M1 protein [8] promotes immune evasion by masking sites for C3 and IgG deposition on the bacterial surface, decreasing the amount of C3 convertase on the bacterial surface, and blocking the receptors for C3bi on macrophages [9]. Furthermore, M protein can assist in adhesion to the extracellular matrix by binding to fibronectin [10] and various types of collagen [11,12].

Beyond its critical role in immune evasion, there is clear evidence that M1 protein also functions as a proinflammatory factor when present as a soluble protein, such as by inducing the release of heparin binding protein from neutrophils, thought to lead to the hypovolaemic shock typical of streptococcal toxic shock [13]. Soluble M1 protein also contributes to NLRP3 inflammasome activation [14] and is capable of activating platelets and interacting with haemostasis [15–17].

In our studies to explore the relationship of *S. pyogenes* with lymph nodes, we identified isogenic strains that were selected for, and enriched within, murine lymph nodes during experimental invasive infection [18]. In all previous studies to date, isogenic strains enriched in lymph nodes have had mutations that result in increased hyaluronan capsule, the ligand for lymphatic endothelial receptor, LYVE-1. In the current study, we report the emergence of an *S. pyogenes* clone in murine lymph node with a nonsense mutation in the *emm1* gene. As a result, this strain was devoid of M1 protein on its surface but secretes a truncated M1 protein directly into the extracellular milieu. The selective emergence of this strain *in vivo*, despite the widely held belief of M protein essentiality, underlines a gap in our understanding of the role M protein may play at different stages and sites of infection.

## Results

### Characterisation of an *emm*1 truncation mutant following intramuscular infection of mice

To investigate bacterial phenotypes associated with lymph node tropism and lymphatic system survival, bacterial colonies cultured from the draining lymph node of mice infected intramuscularly with a single *S. pyogenes emm*1 isolate, H584, were genome sequenced and compared with the non-mucoid parent strain. Five non-mucoid bacterial colonies cultured from the draining lymph node 24 h after intramuscular infection were sequenced and 5/5 contained the same nonsense SNP within the *emm1* gene; this mutation was not detected in colonies cultured from spleen, or thigh muscle, or the inoculum (accession numbers ERS18607827, ERS18607828, ERS18607829, ERS18607830, ERS18607831). This was the only genetic difference compared to the parent strain. The SNP conferred a premature stop codon in *emm1* (GAA to TAA) at residue 199/485, which is within the spacer region of M protein and predicted to result in a truncated M1 protein moiety lacking C and D repeat regions and the LPXTG cell wall anchoring motif (Fig 1 A-B). This isogenic mutant derivative was designated H584_XM_, to indicate an e<u>x</u>tracellular truncated <u>M</u>1 protein. The parent strain and resulting mutant strains did not possess any mutations in the CovRS two-component regulator. Western blot using an anti-M1 hypervariable region (HVR) antibody showed full length M1 protein (54 kDa) in the cell wall extract and supernatant of parent strain H584, but a protein of ∼23 kDa present in the supernatant yet absent in the cell wall extract of mutant strain H584_XM_ (Fig 1C), consistent with the predicted lengths of the surface-associated and secreted M1 protein variants. Strain H584_Δ*emm*1_, an isogenic derivative with allelic replacement of the *emm1* gene (accession number ERS18607832), had no detectable M1 protein in either cell wall extract or supernatant as expected (Fig 1C). To confirm that the nonsense mutation in *emm1* conferred a specific survival advantage in the lymphatic system, the parent strain H584_WT_ and H584_XM_ were compared directly; bacterial survival in different tissues was measured 3 h after intramuscular (i.m) infection. Despite survival in muscle to similar degree, H584_XM_ demonstrated significant survival advantage in the spleen, draining ipsilateral inguinal and axillary lymph nodes compared with the parent strain (Fig 1D).

**Fig 1.**
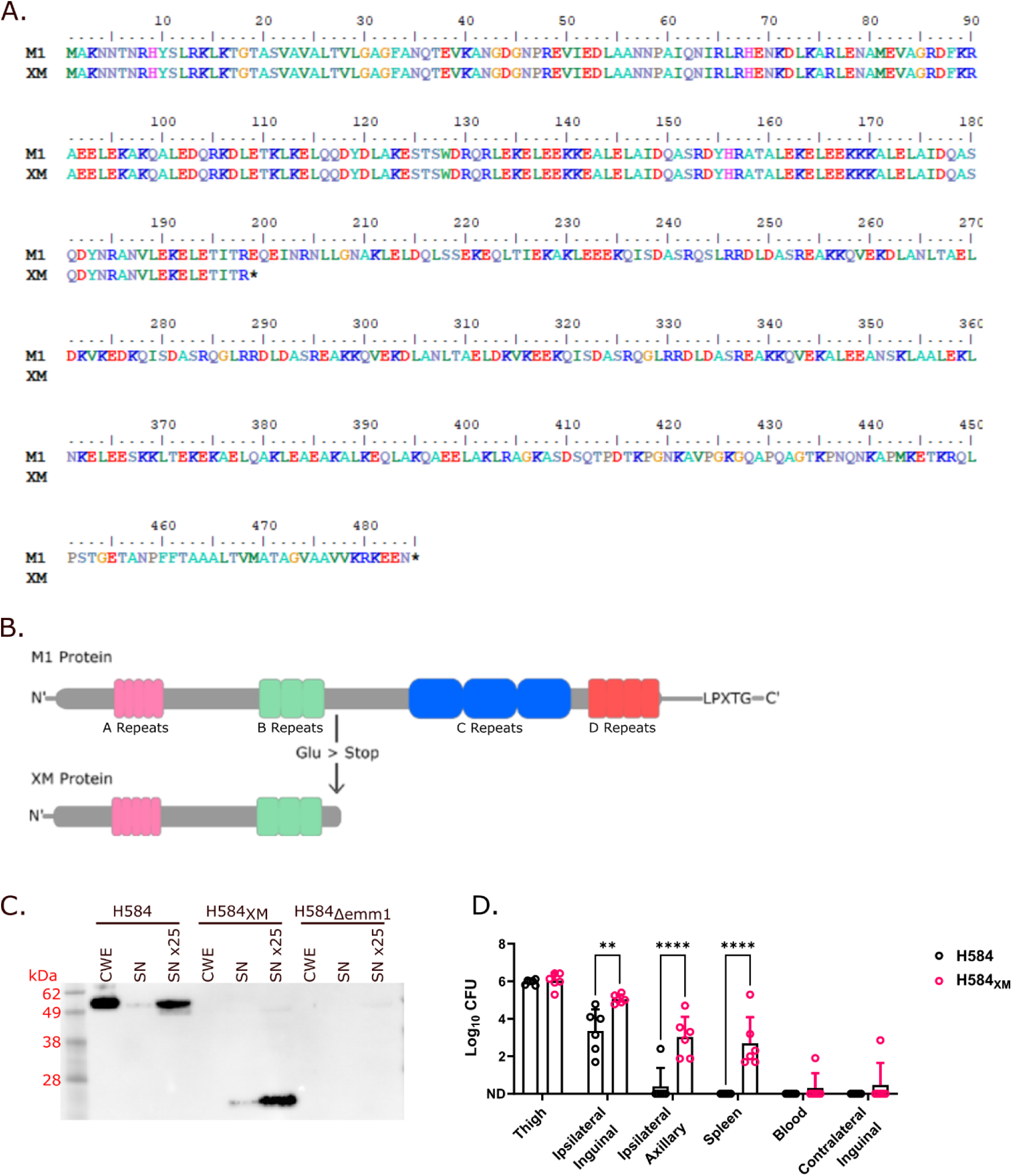
Selective emergence of M1 protein mutant *S. pyogenes* which secretes a truncated M protein. (A) Amino acid alignment showing a stop codon encoded at residue 199 and (B) diagrammatic representation of the XM protein resulting from the SNP in the *emm*1 gene of H584_XM_ compared to wild type M1 protein. (C) H584_XM_ secretes a 23 kDa truncated form of M1 protein (XM protein) directly into the supernatant (SN), with no detectable XM protein found in the cell wall extract (CWE); comparison with parent strain and emm1 deletion mutant. (D) Following i.m. infection, H584_XM_ was recovered in significantly higher numbers from the draining (ipsilateral) lymph nodes and the spleen compared to parent strain H584 3 h after intramuscular infection of FVB/n mice (CFU/mg thigh tissue, CFU/lymph node, CFU/spleen, and CFU/ml of blood) (** p<0.01, **** p<0.0001). ND: not detected.

### Truncating mutation in *emm*1 is associated with both enhanced lymphatic system survival and soft tissue spread in mice

To determine if the expansion of H584_XM_ within the lymphatic niche was specific to the FVB/n mouse strain, that has a known defect in complement factor C5 [19], experiments were repeated in BALB/c mice. To differentiate effects on lymphatic transit from effects on lymphatic system survival, the experiment was undertaken at both an early time point, 3 h, and a later time point, 24 h, after i.m. infection with isogenic strains (H584, H584_XM_ and H584_Δ*emm*1_). Three hours after infection, quantities of streptococci recovered from the draining (ipsilateral) lymph nodes were similar regardless of infecting strain, indicating that, in BALB/c mice, cell-surface M1 protein expression does not affect *S. pyogenes* transit to local lymph nodes during the initial stages of infection (Fig 2A). Notably at this early time point, there was little if any bacterial growth from contralateral lymph nodes, highlighting the role of the ipsilateral lymphatics in *S. pyogenes* transit from the infection site in early infection.

**Fig 2.**
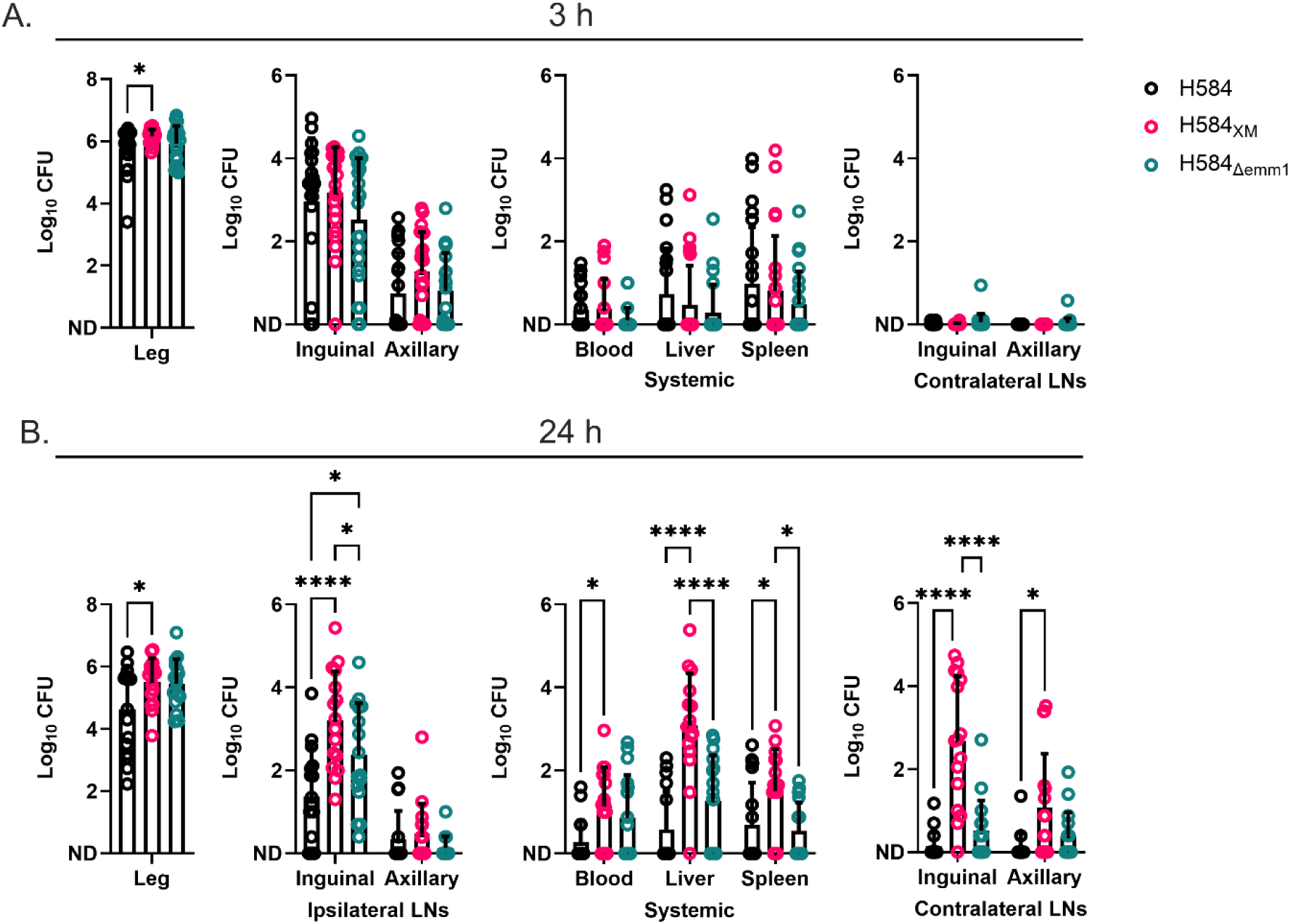
Effect of *emm1* mutation on bacterial survival in lymph nodes and systemic spread at early and late time points. BALB/c mice were infected intramuscularly with 2 × 10^8^ CFU/mouse. (A) Absence of cell wall-associated M protein did not affect bacterial load in the leg (CFU/100 mg tissue), blood (CFU/ml), spleen (CFU/spleen) or liver (CFU/100 mg tissue) and had no effect on bacterial burden in draining (ipsilateral) lymph nodes 3 h post infection (CFU/lymph node) (three pooled experiments, n=22) (B) At 24 h post infection H584_XM_ exhibited significantly greater ipsilateral and contralateral lymph node survival and systemic spread (two pooled experiments, n=16). (* p<0.05, **** p<0.0001). ND: not detected.

In contrast to the observations made 3 h post infection, 24 h after i.m. *S. pyogenes* infection, both mutant strains lacking M protein on the cell wall were found in significantly higher numbers in the draining lymph node (ipsilateral inguinal) than the parent strain H584 (Fig 2B).

Indeed, H584_XM_ was recovered at higher numbers than H584 in ipsilateral inguinal lymph node, blood, liver, and spleen, despite the inoculum being similar for all groups. Infection with H584_XM_ caused greater systemic spread than both wildtype strain H584 and H584_Δ*emm*1_ (Fig 4B). This difference could not be explained by a simple growth advantage as growth in broth (as measured by CFU) was similar for all three strains (S1 Fig).

Interestingly, mice infected with *S. pyogenes* H584_XM_ exhibited significantly greater bacterial loads within the contralateral inguinal lymph node by 24 h compared to mice infected with other strains. The contralateral inguinal lymph node does not usually drain the site of infection used in this model [20]. Although dissemination of bacteria to contralateral lymph nodes can occur due to direct seeding via the systemic blood circulation, the *S. pyogenes* H584_XM_ load in the contralateral inguinal lymph node was an order of magnitude greater than measured in the contralateral axillary lymph node. This is consistent with spread of the soft tissue infection across the midline of the mouse, with consequent lymphatic drainage to the opposite inguinal lymph node. Consistent with this, histopathological examination of infected thigh muscle and skin, overlying abdominal muscle and skin, and the pelvic girdle of infected mice demonstrated infection with H584_XM_ to be associated with increased inflammation, tissue necrosis and bacterial dissemination in the abdominal and pelvic wall tissues compared with mice infected with parent strain H584. This was despite the site of infection in thigh muscle appearing similar, regardless of which strain had been used for infection (Fig 3A) (full summary in S1 Table). The results suggest that H584_XM_ not only exhibits a survival advantage in the lymphatic system but is associated with increased local bacterial dissemination and more widespread tissue damage following soft tissue infection, leading to drainage into non-standard lymph nodes, contralateral to the site of infection (Fig 3B).

**Fig 3.**
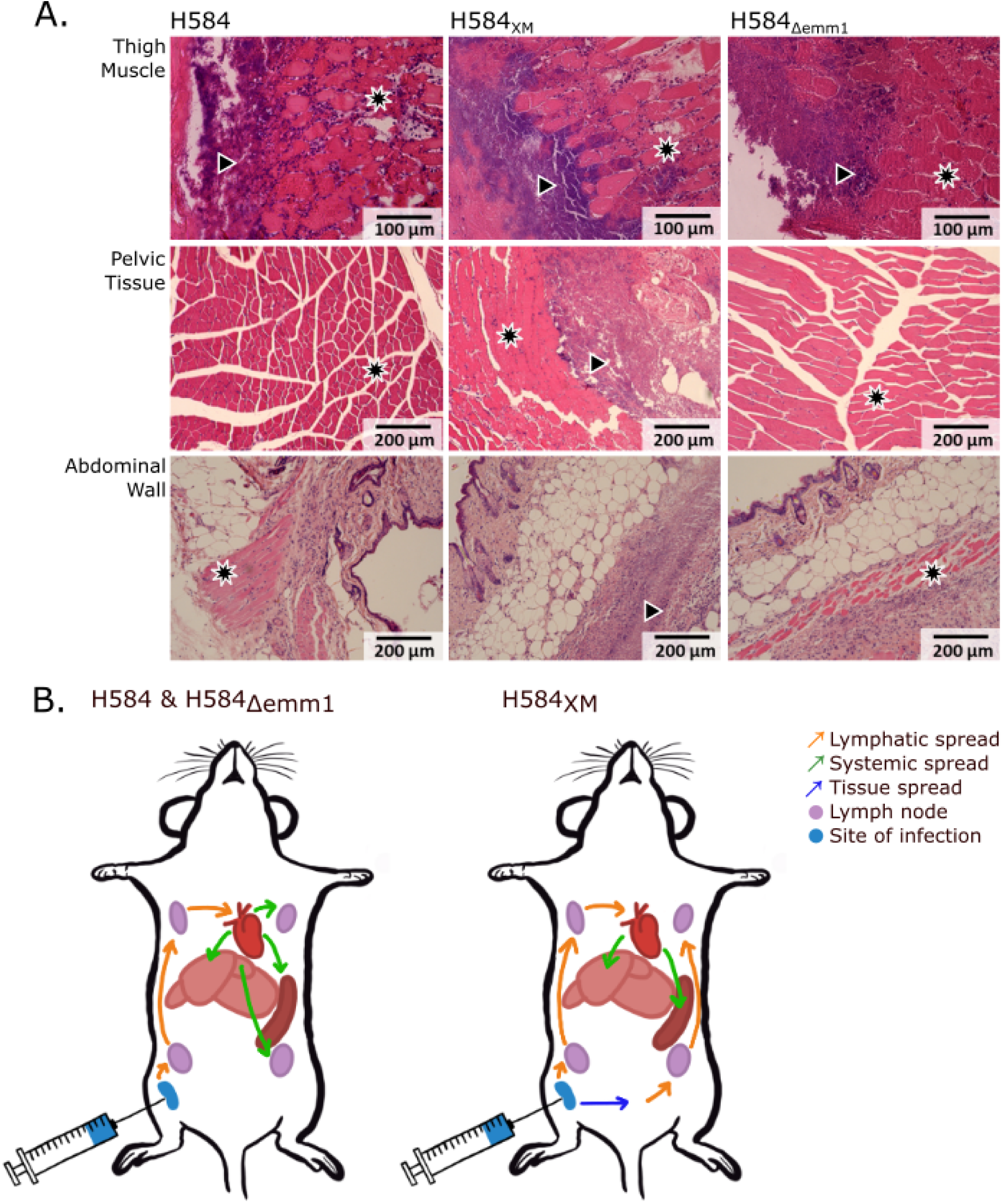
H584_XM_ infection exhibits a different dissemination pattern to H584 and H584_Δemm1_. (A) Histopathological examination of H&E-stained thigh muscle, pelvic floor tissue, and abdominal wall tissue from mice infected intramuscularly for 24 h showed increased tissue necrosis, inflammation and bacteria present in abdominal and pelvic tissue from mice infected with H584_XM_. Healthy tissue appears pink in colour (indicated by star symbol), while bacterial infiltrate and inflammation appear purple (indicated by arrowhead symbol). Images representative of four mice per group. (B) Diagram showing inferred dissemination routes via lymphatic system and systemic circulation of strains at 24 h post infection. H584_XM_ caused lateral spread of infection associated with increased spread to the contralateral lymphatic system.

To further isolate the effects of M protein on bacterial lymphatic transit, we infected mice with *Lactococcus lactis* heterologously expressing either full-length cell-wall associated M protein or secreted XM (S2 Fig). In contrast to *S. pyogenes*, cell wall-associated M protein conferred a survival advantage to *L. lactis* in thigh muscle. Although there were insufficient bacteria in the draining lymph node at the 3 h time point to make a comparison, 24 h post-infection, the survival of *L. lactis* expressing XM in the draining lymph node was equivalent to *L. lactis* expressing full length M protein, despite a log_10_ fold reduction in bacteria expressing XM in the infected thigh tissue.

### H584_XM_ has a competitive advantage in experimental soft tissue infection

As H584_XM_ had emerged following infection with a parent *emm*1 strain that expressed full length M protein, we investigated the likelihood that such a mutant would outcompete the parent strain. The ability of H584_XM_ to compete against H584 was determined using intramuscular infection with a mixed inoculum. Despite being infected with a 1:1 mixture of H584 to H584_XM_, after 3 hours of infection, H584_XM_ exhibited a 4-fold advantage relative to H584 in the draining lymph node, in contrast to the site of infection where the ratio was similar to the original inoculum (Fig 4A). The progression of H584_XM_ infection compared to H584 was not replicated by infection with H584_Δ*emm*1_ pointing to a role for the secreted XM in conferring favourable advantage. The competitive advantage of H584_XM_ was further augmented 24 h post infection, in both the infected hind limb and the draining inguinal lymph node (Fig 4B).

**Fig 4.**
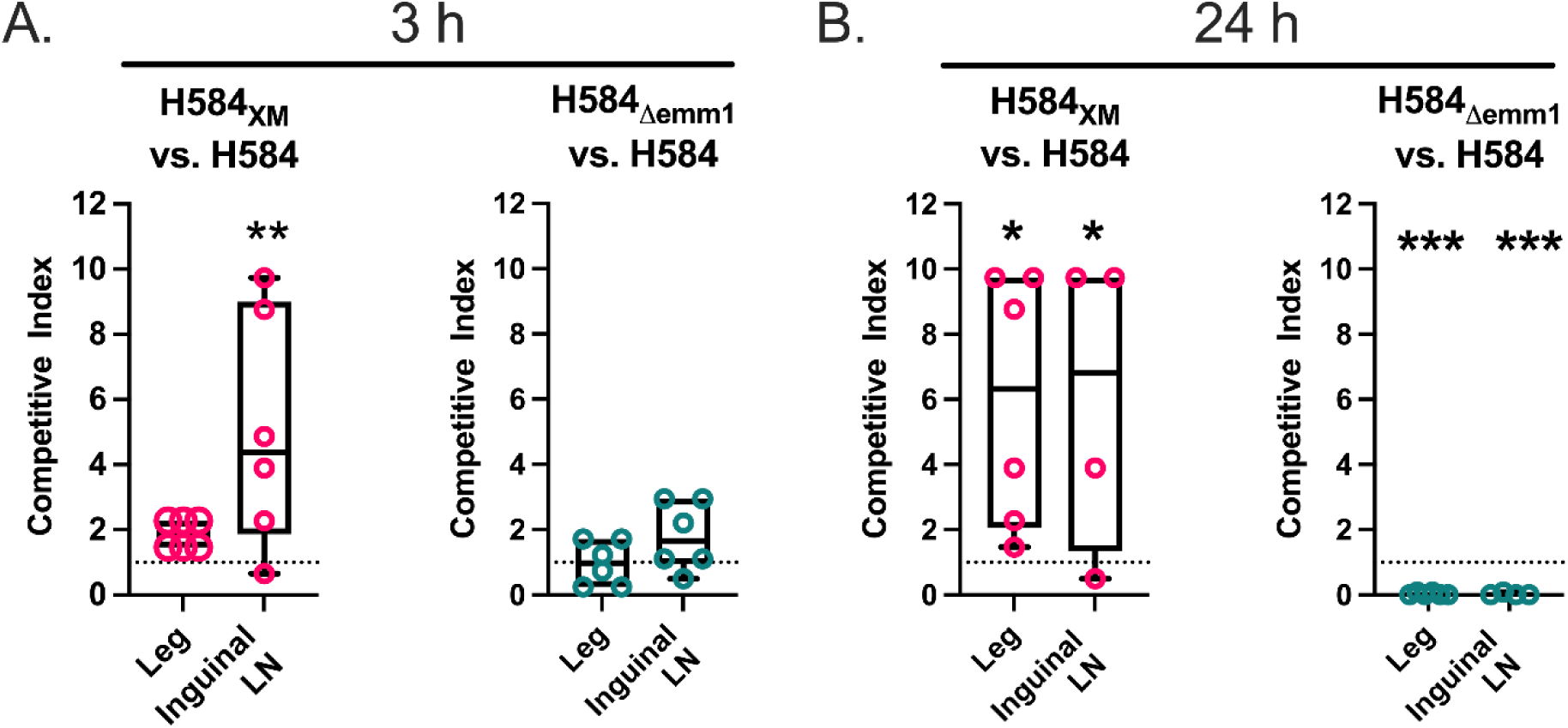
H584_XM_ outcompetes H584 in soft tissue infection. Following intramuscular infection of BALB/c mice with an inoculum composed of 1:1 mixtures of strains, H584_XM_ outcompeted H584 in the leg and ipsilateral draining lymph node at 3h (A) and 24 h (B) post infection while H584_Δemm1_ had no competitive advantage over the WT strain. Each point represents an individual mouse and indicates the competitive index (C.I.), which is the final ratio of the mutant strain to the parent strain H584 at the indicated time point (C.I.>1 indicates superiority). Statistical analysis compared final ratio with initial ratio in the inoculum (* p<0.05, ** p<0.01, *** p<0.001). Dotted line represents ratio of inoculum.

Taken together with the earlier experiments, the data showed that in the initial stages of soft tissue invasive infection, M protein is not required for infection progression and, furthermore, *S. pyogenes* strains that shed M protein may have an advantage as invasive soft tissue infection progresses.

### Cell surface M protein is required for survival in whole human blood, but not whole mouse blood

*S. pyogenes* cell surface M protein is recognised to be important for resistance to opsonophagocytosis in human blood, although H584_XM,_ that lacks surface M protein, was able to survive and outcompete wild type H584 in mice. To determine if the isogenic strains exhibited the anticipated survival phenotypes in whole blood, H584, H584_XM_ and H584_Δ*emm*1_ were co-incubated in heparinized whole human blood and the ability of each strain to multiply over 3 h was assessed. As expected, expression of M1 protein on the surface of *S. pyogenes* was required to permit net growth in all donor blood samples, while growth of both *emm*1 mutants was more limited (Fig 5A). Interestingly, when the experiment was repeated in murine whole blood, this difference disappeared; indeed, H584_XM_ displayed greater multiplication in murine blood than both H584 and H584_Δ*emm*1_ (Fig 5B).

**Fig 5.**
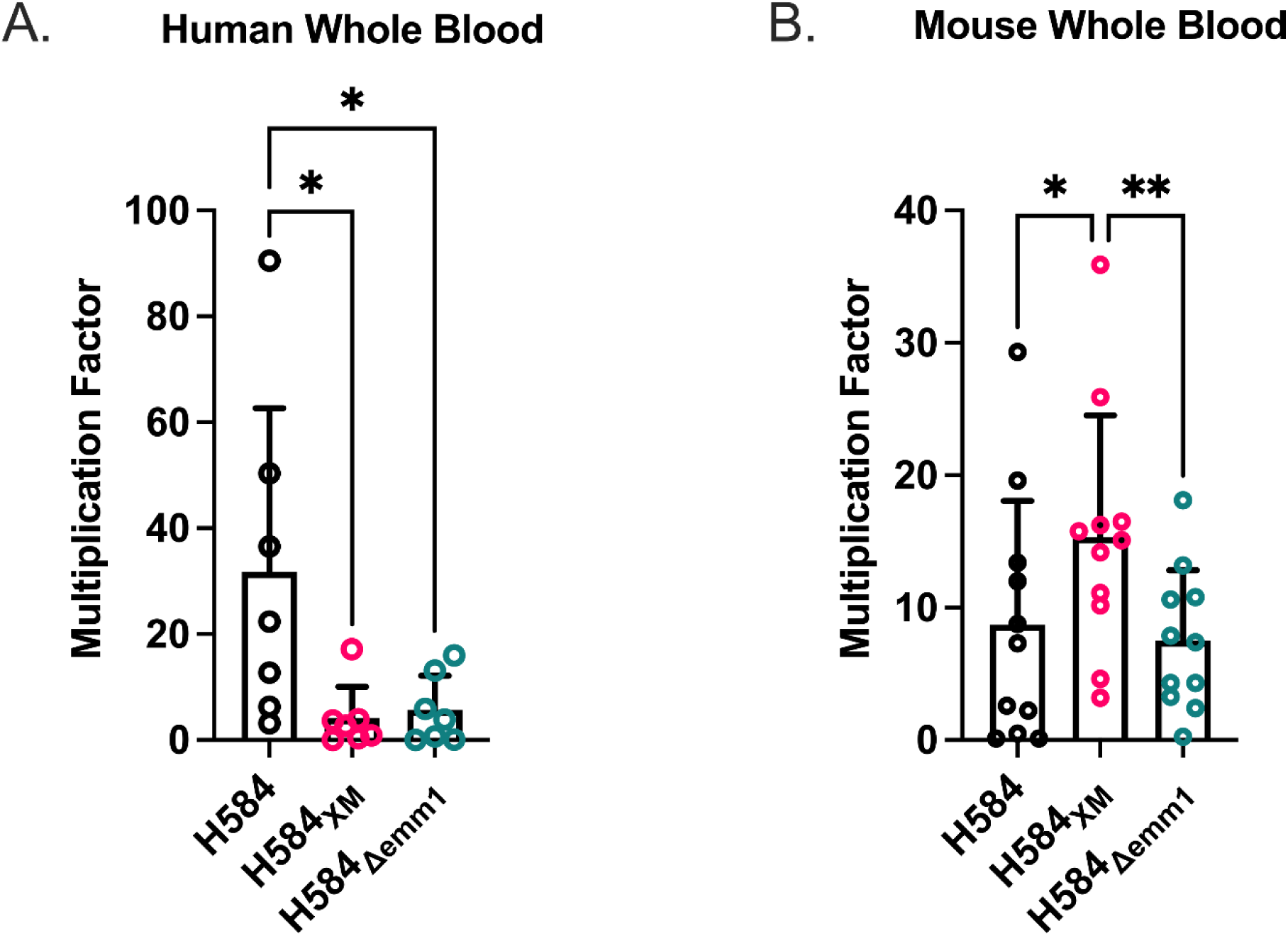
Survival of isogenic M1 strains in whole human blood and whole mouse blood. (A) Expression of M1 protein in the bacterial cell wall augments growth and survival in whole human blood (n=7 individual donors, mean of 3 replicates for each donor shown). (B) Despite lack of cell surface M protein, growth and survival of H584_XM_ was increased compared with H584 and H584_Δemm1_ in whole mouse blood (n=11 individual BALB/c mice, female) (* p<0.05, ** p<0.01). Mean and SD shown.

### Cell wall associated M1 protein is necessary to establish nasopharyngeal carriage of *S. pyogenes*

We considered the possibility that the mutation in H584_XM_ had an unforeseen wider impact on virulence *in vivo* that resulted in generalised enhanced virulence, and so investigated the ability of the same isogenic strains to establish infection in a well-characterised mouse model of nasopharyngeal infection, in which streptococci persist in nasal turbinates for 5–7 days [21]. In contrast to findings in soft tissue infection, cell wall expressed M1 protein was required to sustain nasal carriage of *S. pyogenes* in FVB/n mice. Mice infected intranasally with H584_XM_ or H584_Δ*emm*1_ cleared bacteria from the nasopharynx significantly faster than mice infected with parent strain H584, and these mice also had decreased airborne shedding of bacteria (Fig 6A-C). When infected with a similar number of CFU in a greater volume (10 µL), a model of intranasal infection which is more likely to lead to lower respiratory tract infection and systemic disease [22], all mice infected with H584 progressed to systemic infection within 72 h, whereas this was seen in only a minority of animals infected with H584_XM_ or H584_Δ*emm*1_ (Fig 6D). The bacterial burden in the nasopharynx of mice infected with the wildtype strain H584 exceeded that of mice infected with H584_XM_ or H584_Δ*emm*1_ by two orders of magnitude. This difference was amplified even further when considering bacterial dissemination to liver and spleen (Fig 6D).

**Fig 6.**
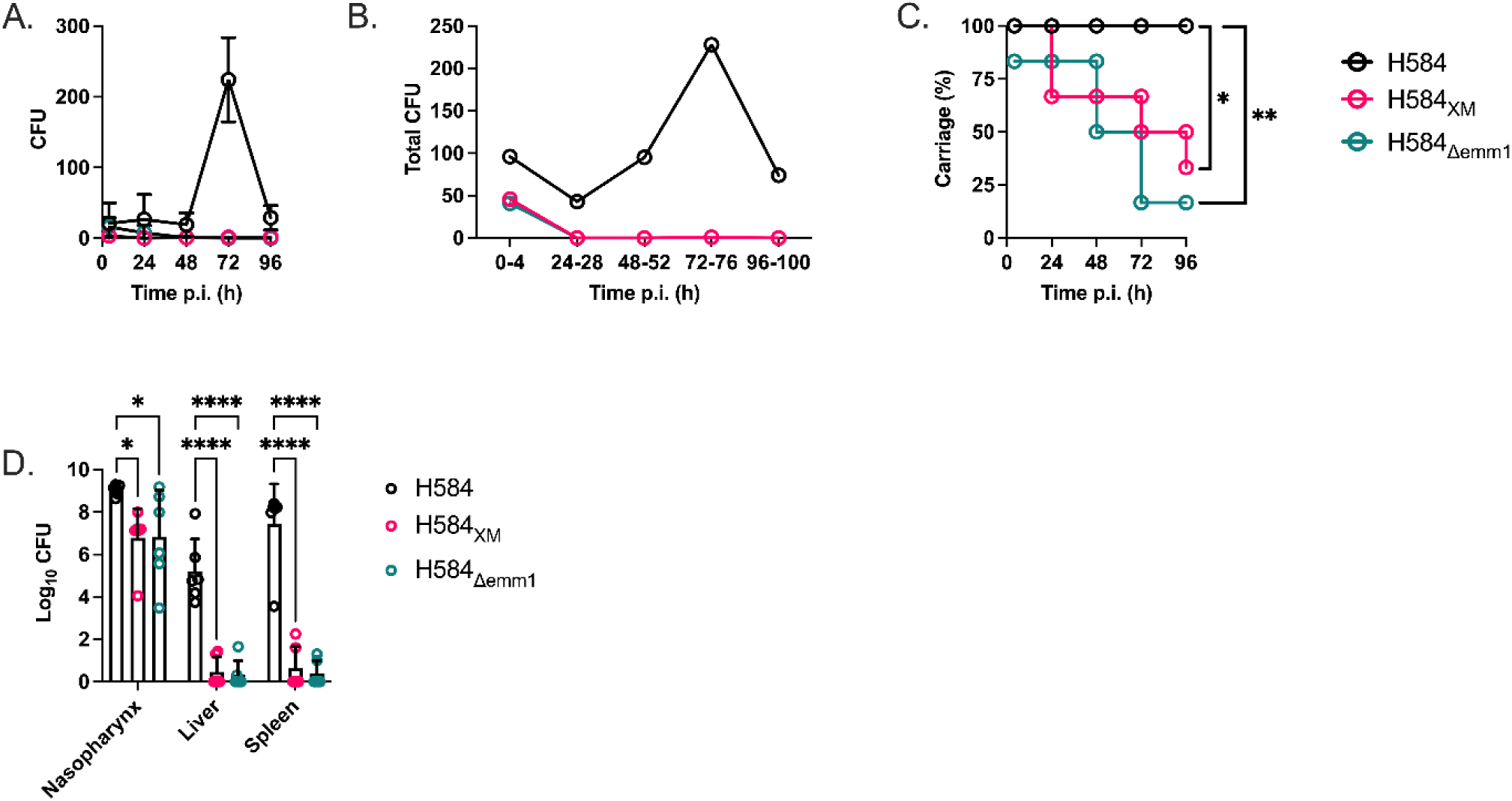
Cell wall associated M1 protein is required to establish nasopharyngeal infection in FVB/n mice and necessary for progression to systemic infection via the lower respiratory tract route. (A) Following nasal infection, mice infected with wildtype strain H584 demonstrated heavier nasal shedding of *S. pyogenes* than those infected with H584_XM_ and H584_Δ*emm1*_ (mean and SD shown). (B) Mice infected with H584 shed greater amounts of bacteria into the air as determined by cage settle plates over the course of 4 days. (C) Nasopharyngeal clearance of *S. pyogenes* occurred significantly earlier in mice infected with H584_XM_ or H584_Δ*emm1*_ compared to H584. (D) Following intranasal infection with a high dose in a large volume, only wildtype strain H584 was reproducibly recovered in high numbers from liver and spleen (CFU per whole nasopharynx and spleen, CFU per 100 mg liver shown) (n=6, * p<0.05, ** p<0.01).

### M1 protein expression modulates binding to extracellular matrix protein and glycans components

We considered the possibility that secretion of M1 protein, and consequent loss of cell wall associated M1 protein, resulted in unforeseen differences in streptococcal adhesion to host factors that might explain lymphatic tropism or enhanced survival.

Recombinantly produced XM protein, (designed to emulate the secreted truncated M protein produced by H584_XM_) was able to bind fibrinogen and fibronectin to the same extent as recombinantly produced full length M1 protein (Fig 7A). XM protein showed a reduced ability to bind collagen types I and IV, which was expected, as the region of M1 protein that mediates collagen binding is missing from XM (Fig 7A).

**Fig 7.**
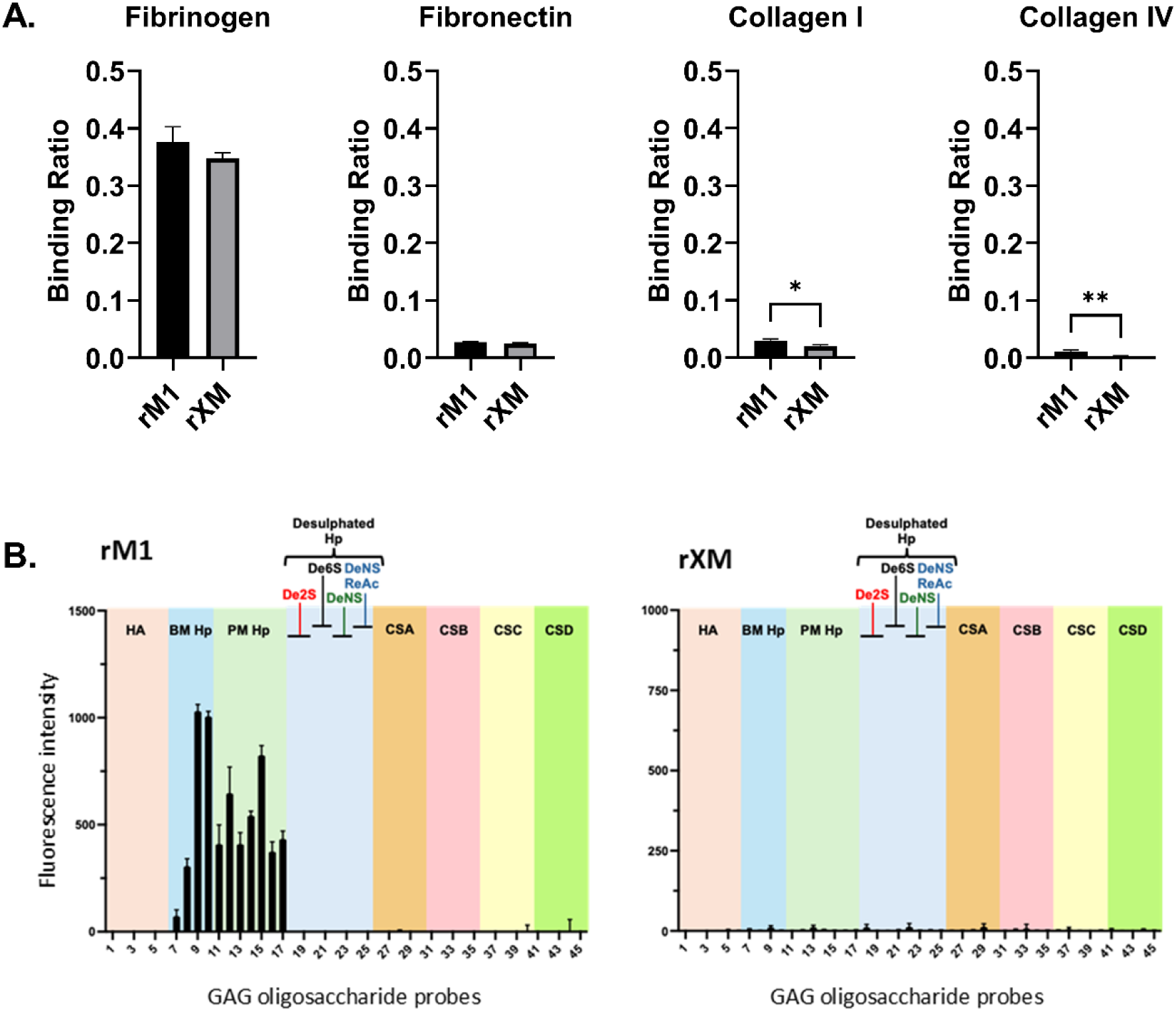
M protein is an important mediator of *S. pyogenes* interactions with host proteins and glycosaminoglycans (GAGs) (A) Recombinant XM protein (rXM) can bind to fibrinogen and fibronectin at similar levels to full length M1 protein (rM1) but is unable to bind collagen types I and IV (data represent the mean +/− the standard deviation of three replicates; * p<0.05, ** p<0.01, *** p<0.001, **** p<0.0001). (B) Mean fluorescence intensities of His-tagged rM1 and rXM proteins binding to 45 oligosaccharide probes on a focused GAG oligosaccharide microarray (S1 File; S2 Table). rM1 shows selective binding to highly sulfated heparin (Hp) probes derived from bovine mucosa (BM) and porcine mucosa (PM), but not to site-specific desulfated Hp or probes representing other GAG classes. In contrast, rXM shows no detectable binding to any of the GAG oligosaccharides tested. Different GAG classes are indicated by the coloured panels as defined at the top of the histogram charts: hyaluronic acid (HA), heparin (Hp), site-specific desulfated Hp, and chondroitin sulfate (CS) A, B, C, and D. Data are presented as background-subtracted mean fluorescence intensities of quadruplicate spots for each oligosaccharide probe; error bars represent the standard deviation (SD).

Glycosaminoglycans (GAGs) such as chondroitin sulfate, dermatan sulfate, heparin/heparan sulfate and hyaluronan are highly abundant within the mammalian extracellular matrix and at the cell surface. We therefore evaluated the glycan-binding specificities of rM1 and rXM proteins using a broad-spectrum neoglycolipid (NGL)-based glycan screening microarray containing 672 lipid-linked glycan probes [23](S3 Fig) and a focused GAG oligosaccharide microarray (Fig 7B). Notably, rM1 bound exclusively to heparin-related oligosaccharide probes in both broad-spectrum screening array and the focused GAG array, with signal intensity increasing alongside chain length (S1 File). No binding was detected to HA (non-sulfated GAG), desulfated heparin or CS probes (Fig 7B), highlighting the importance of degree of sulfation for M1-GAG interaction. In contrast to the strong heparin binding with M1, the truncated XM protein showed no detectable binding to short heparin oligosaccharides (<DP16) in the focused GAG array (Fig 7B), although binding to longer heparin chains (DP18-DP20) was detected in the NGL-based glycan screening microarray (S3 Fig; S1 File), indicating a potential preference of XM for long heparin chains.

As expected, *S. pyogenes* strain H584 was able to bind fibrinogen, fibronectin, and collagen types I and IV to a significantly greater degree than H584_XM_ and H584_Δ*emm*1_ (Fig 8A). No differences were observed between H584_XM_ and H584_Δ*emm*1_ in any of the adhesion assays. To determine if the isogenic streptococcal strains differed in glycan binding, fluorescently labelled H584, H584*_XM_* and H584_Δ*emm1*_ were tested for GAG binding on a GAG polysaccharide microarray containing CSA, CSB, CSC, heparin (a commonly used model for the highly sulfated domains of heparan sulfate), and hyaluronan polysaccharides (Fig 8B) (S1 File). The parent strain H584 showed binding to heparin polysaccharide, which was reduced in the H584_Δ*emm*1_ strain (Fig 8B and S4 Fig). In contrast, an unexpected increase in binding to hyaluronan was demonstrated by both H584_Δ*emm*1_ and H584*_XM_*. Chondroitin sulphates were not bound by any strain (S1 File and S4 Fig).

**Fig 8.**
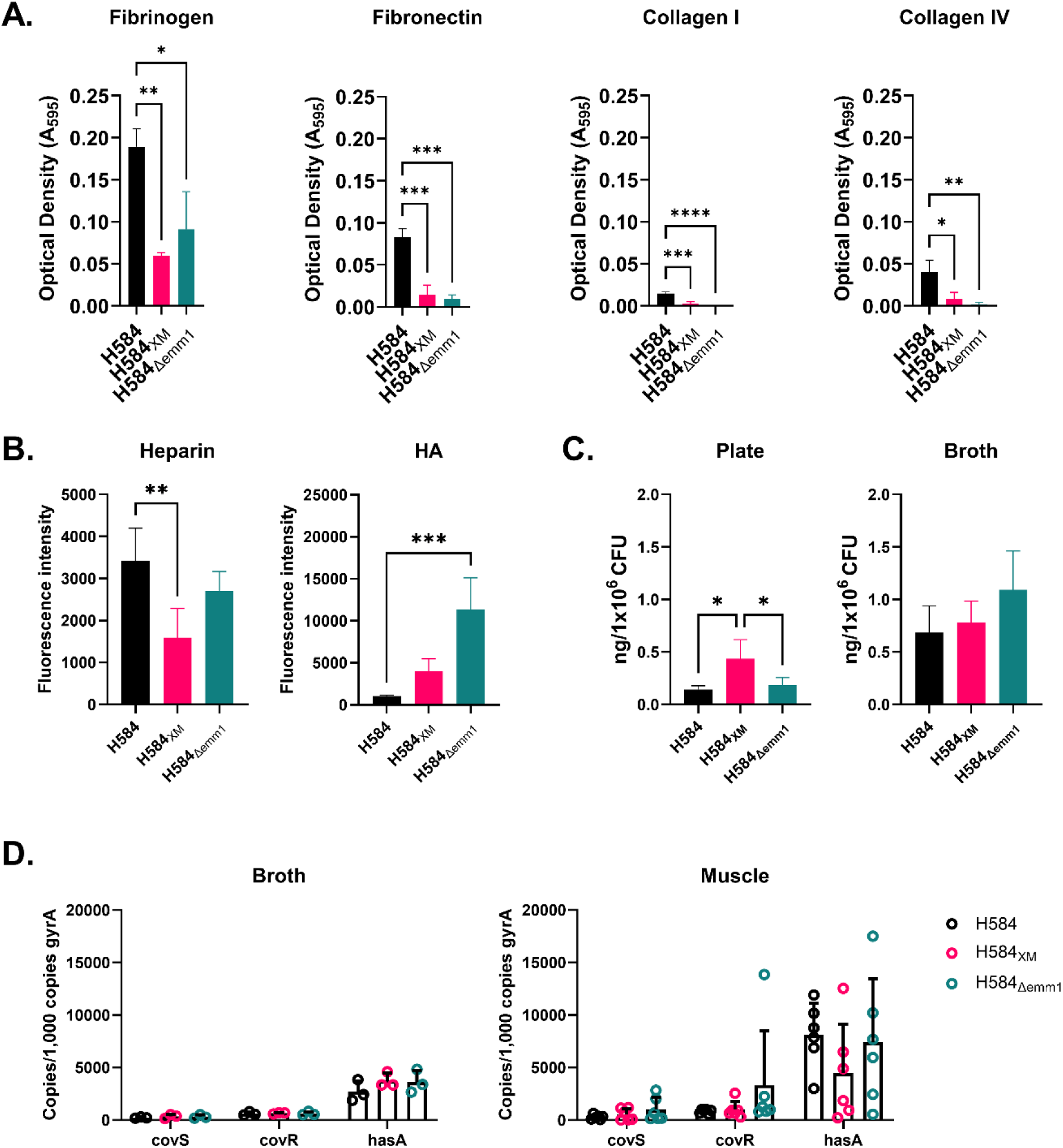
GAG binding is modulated by M1 protein expression. (A) Loss of cell wall associated M protein significantly affects binding of *S. pyogenes* to fibrinogen, fibronectin, collagen type I and collagen type IV (data represent the mean +/− the standard deviation of three biological replicates) (B) Mean fluorescence intensities of fluorescently labelled H584, H584_XM_, and H584_Δemm1_ binding to biotinylated GAG polysaccharides printed at 0.5 mg/ml (Hyaluronan) or 0.1 mg/ml (Heparin). The background subtracted mean fluorescence intensities of quadruplicate spots on the microarray is shown. Error bars represent SD of quadruplicate spots for each polysaccharide. Assay is representative of n=3. Full GAG polysaccharide array data are shown in S4 Fig and S1 File. (C) The amount of hyaluronan capsule expressed by the panel of strains was measured from bacteria grown overnight on CBA (n= 3 biological replicates) and mid-log bacteria grown in THB (n=3 biological replicates); only H584XM grown on CBA showed increased HA compared to the other strains (p<0.05). (D) Transcription of the genes *covR*, *covS* and *hasA* from *S. pyogenes* grown in THB (n=3) or within thigh tissue of mice 24 h after intramuscular infection (n=6) or was measured using qRT-PCR. No difference in transcription between the strains was observed. Transcription was compared to that of housekeeping gene *gyrA*. (* p<0.05, ** p<0.01, *** p<0.001, **** p<0.0001.)

As the isogenic M protein mutants were more able to bind exogenous hyaluronan, we considered whether H584_XM_ and H584_Δ*emm*1_ were better able to retain capsular hyaluronan, thus increasing the amount of capsule in a manner independent of *has* operon transcription. When hyaluronan was extracted from mid logarithmic phase bacteria grown in liquid culture, no significant differences were seen (Fig 8C) but when hyaluronan was extracted directly from bacteria grown on solid media H584_XM_ had significantly increased hyaluronan (Fig 8C). We did not identify differences between isogenic strains in transcription of the HA synthesis gene *hasA,* or the genes of capsule regulator, *covS and covR,* in bacterial RNA recovered from infected thigh tissue, and there was no difference in transcription of these genes when bacteria were grown in standard media (Fig 8D).

## Discussion

M protein is historically one of the most investigated virulence factors of *S. pyogenes* [7,24–29]. In this work we have identified and characterised a strain of *S. pyogenes* that exclusively secretes a soluble form of M1 protein into the extracellular milieu. This XM-secreting derivative resulted from just a single SNP and exhibited enhanced survival in both the lymphatic system and soft tissues, when injected directly. The phenotype of the XM mutant was only partially recapitulated by the strain lacking any M protein, demonstrating that the distinctive observations attributed to H584_XM_ are not solely due to the absence of cell wall anchored M1 protein but are also mediated by the expression of soluble XM protein. The study has shown an almost absolute requirement for *S. pyogenes* M1 protein surface expression in acquisition and persistence of murine nasopharyngeal infection, and dissemination from the lung, but no requirement for M1 protein surface expression during invasive soft tissue infection, once infection is initiated. The unexpected expansion and apparent survival advantage of an *S. pyogenes* strain without cell wall anchored M1 protein demonstrate that there is much yet to learn about the role of M protein during different types of experimental streptococcal infection.

*In vivo* infections revealed two specific and unexpected phenotypes of the XM mutant. Firstly, H584_XM_ showed increased survival specifically in draining lymph nodes in comparison with the parent strain. Such lymph node tropism has previously been associated with mutations that lead to increased hyaluronan capsule expression, either via *covRS* or *rocA* regulatory gene changes, or by alterations of the capsule promoter [18], leading to interaction with the lymphatic endothelial receptor LYVE-1[30]. However, the isolates in this study selected in lymph nodes had no mutations in the capsule locus and transcriptional studies revealed no changes in transcription of genes associated with capsule. Importantly, glycan binding studies confirmed that isogenic strains H584_XM and_ H584_Δ*emm*1_ bound more exogenous hyaluronan than the parent strain. Furthermore, there was evidence that H584_XM_ retained more of its endogenous HA capsule when cultured on solid media than the parent strain. Taken together we construe that the absence of cell surface M protein leads to enhanced coating of H584_XM in_ host hyaluronan as well as improved retention of streptococcal capsule. The nature of the streptococcal ligands that engage hyaluronan are at present unknown, but increased surface hyaluronan, whether of mammalian or bacterial origin, provides an explanation for lymph node tropism.

Secondly, the H584_XM_ isolate showed a generalised increased competitive advantage compared with the parent strain and was associated with increased inflammation and spread during soft tissue infection. This was perplexing given the dogma that M protein is required for resistance to infection particularly in blood. The *in vitro* whole blood assays suggest that there may be a human-specific role for M protein in resistance to phagocytosis, that is not seen in mouse blood, or that pre-existing antibodies present in adult human blood may be required for the effects of M protein. This could be of importance when considering *S. pyogenes* pathogenesis in young non-immune children or in niches that lack immunoglobulin. The spread of infection was specific to H584_XM_ and was not seen in the M protein-negative mutant. We believe that the well-documented pro-inflammatory effects of soluble M1 protein go some way to explain the phenotype observed.

H584_XM_ demonstrated an infection phenotype beyond that of the M protein deletion strain, pointing to further virulence conferred by secretion of XM protein. Soluble M1 protein is reported to promote inflammation through several mechanisms [13–15,31,32], often involving binding to fibrinogen. Furthermore, M1 protein has been implicated in supporting *S. pyogenes* escape from the lysosome, so XM protein expression could provide another advantage, allowing increased intracellular survival [33–35]. We speculate that the presence of greater numbers of H584_XM_ within the contralateral (i.e. non-draining) inguinal lymph node 24 hours after intramuscular infection could be explained by increased tissue damage at the site of infection, thus facilitating lateral dissemination of the bacterial infection, causing drainage to those lymph nodes.

The requirement for M1 protein on the streptococcal surface is evidently site-specific, as in the current study M1 protein was essential to gain a foothold in the murine nasopharynx. We believe this is due to the adhesive properties of M1 protein [33], that are critical in the murine nasopharynx but may be less critical during an inoculation-initiated invasive infection, or an infection that spreads to the lymphatic system. Interpretation of studies examining M protein in rodent nasopharyngeal infection [33,36,37] is complicated by the potential for unveiling alternative ligands in M protein mutants. In combination with the results presented in the current work, these studies point to a role for M protein in promoting persistence in the nasopharynx beyond the initial infection event and early stages of colonisation, at least for *emm1 S. pyogenes*.

Strain H584_XM_ has highlighted the distinct roles played by M protein that are dependent on infection context. The lack of anchored M1 protein that is common to both H584_XM_ and H584_Δ*emm*1_ illustrates a key role for cell wall anchored M1 protein in nasopharyngeal infection, but not for soft tissue infection, nor systemic infection, at least in mice. These findings underline the importance of considering each infection niche when investigating pathogens that are not restricted to a single niche of infection.

The nonsense mutation leading to H584_XM_ we speculate arose during infection. The SNP could not be identified in the original bacterial stock, nor did it arise again in repeat experiments using the same stock. Nonetheless, there is a risk that similar mutations could arise in already established infections and expand thereafter. There is scope for M protein variation to arise without adversely affecting the overall fitness of *S. pyogenes* [38–41]; indeed, this variation could be beneficial to the ability of *S. pyogenes* to further evade the host immune response. In a situation where there is increased evolutionary pressure on M protein, for example through vaccination against M protein, the findings in this study demonstrate how a single mutation could undermine such protection, and in extreme cases lead to the emergence of highly invasive clones.

## Methods

### Ethics statement

Human blood was obtained from consenting healthy donors from an approved subcollection of the Imperial College NHS Trust Tissue Bank (ICHTB reference R12023; HTA licence 12275). In vivo experiments were performed in accordance with the Animals (Scientific Procedures) Act 1986 and were approved by the Imperial College Ethical Review Process (ERP) panel and the UK Home Office.

### Bacterial Culture

Bacterial strains used are listed in Table 1. *S. pyogenes* strains were grown in Todd-Hewitt broth (THB) (Oxoid) or on Columbia blood agar with horse blood (CBA) (E and O Laboratories), at 37°C with 5% CO_2_. *Lactococcus lactis* was grown in GM17 broth or agar (Oxoid) at 30°C in O_2_. Media was supplemented with 500 µg/ml kanamycin (Sigma) for *S. pyogenes*, and 10 µg/ml erythromycin (Sigma) for *L. lactis* as appropriate.

**Table 1.** Bacterial strains used, including species and relevant details.

| Strain | Species | Relevant detail | Source |
| --- | --- | --- | --- |
| H584 | *S. pyogenes* | Clinical M1 isolate, postpartum sepsis | [42] |
| H584<sub>xM</sub> | *S. pyogenes* | Animal passage, truncated extracellular M1 protein | This study |
| H584<sub>Δemm1</sub> | *S. pyogenes* | Chromosomal deletion of *emm1* | This study |
| MG1363 | *L. lactis* |  | Kind gift P. Moreillon |
| H1186 | *L. lactis* | MG1363 with empty pOri23 vector | Kind gift P. Moreillon |
| H1187 | *L. lactis* | MG1363 with pOri23M1 | This study |
| H1472 | *L. lactis* | MG1363 with pOri23XM-His | This study |
| H1511 | *L. lactis* | MG1363 with pOriM1-His | This study |
| H1470 | *L. lactis* | MG1363 with pOri23XM | This study |

### Identification of M1 protein mutant strain

H584_XM_ was identified in the draining inguinal lymph node 24 h after soft tissue infection with H584 following antecedent contusion injury as previously published [18]. Non-mucoid colonies from the local draining lymph node were whole genome sequenced using Illumina MiSeq (MRC CSC Core Genomics Laboratory, Imperial College London). Short read sequences were mapped using SMALT (http://www.sanger.ac.uk/resources/software/smalt/) to the reference genome MGAS5005 (Genbank accession number NC_007297.2). The SNP in *emm*1 was manually checked against the whole genome sequence of H584 using Artemis [43]. Whole genome sequences are available on the European Nucleotide Archive (ENA) under project code PRJEB74316.

### Deletion of *emm*1 by homologous recombination

Genomic DNA from *S. pyogenes* H584 was extracted as previously described [44]. To replace the *emm*1 allele, a 536 bp fragment containing the intergenic region between *emm1* and *mga* including 315 bp of the *mga* gene was amplified and cloned into the suicide vector pUCMUT using restriction sites Kpn1 and EcoRI, at the 3’ and 5’ ends, respectively. A second 535 bp fragment containing the intergenic region between *sic*1 and *emm*1 including 335 bp of the *emm*1 gene was amplified and cloned into the plasmid using restriction sites SalI (3’) and PstI (5’), to produce pUCMUT*_emm_*_1KO_. All primers are listed in Table 2. The resulting plasmid was transformed into H584 by electroporation as described [45] and successful transformants selected on kanamycin; the *aphA3* kanamycin resistance gene was introduced into the chromosome by homologous recombination, as confirmed by PCR and sequencing of the region across *sic1*, *emm1* and *mga*. The genome of the resulting strain, H584_Δ*emm*1_, was sequenced as above and compared to the genome of parent strain H584.

**Table 2.**
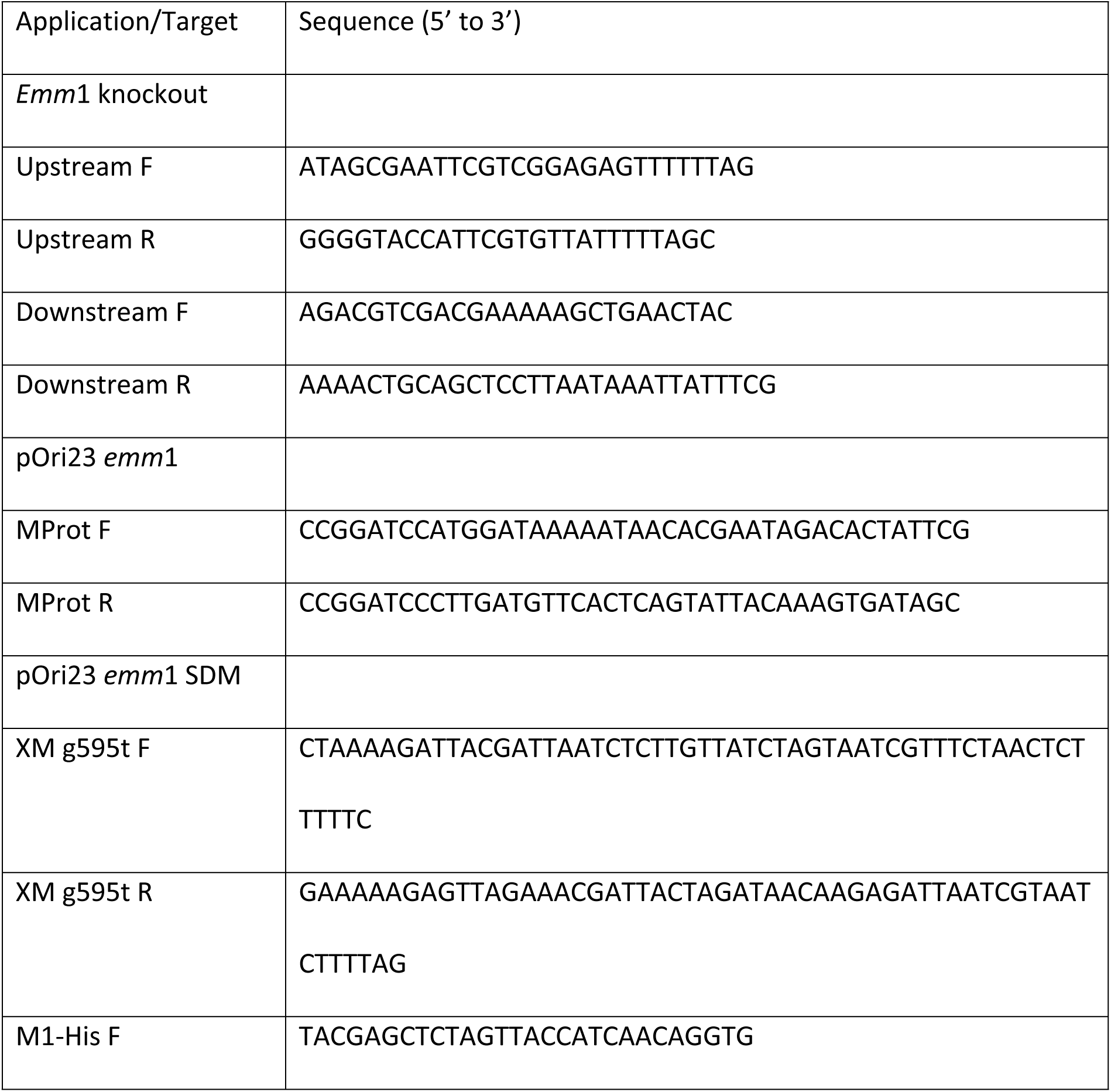
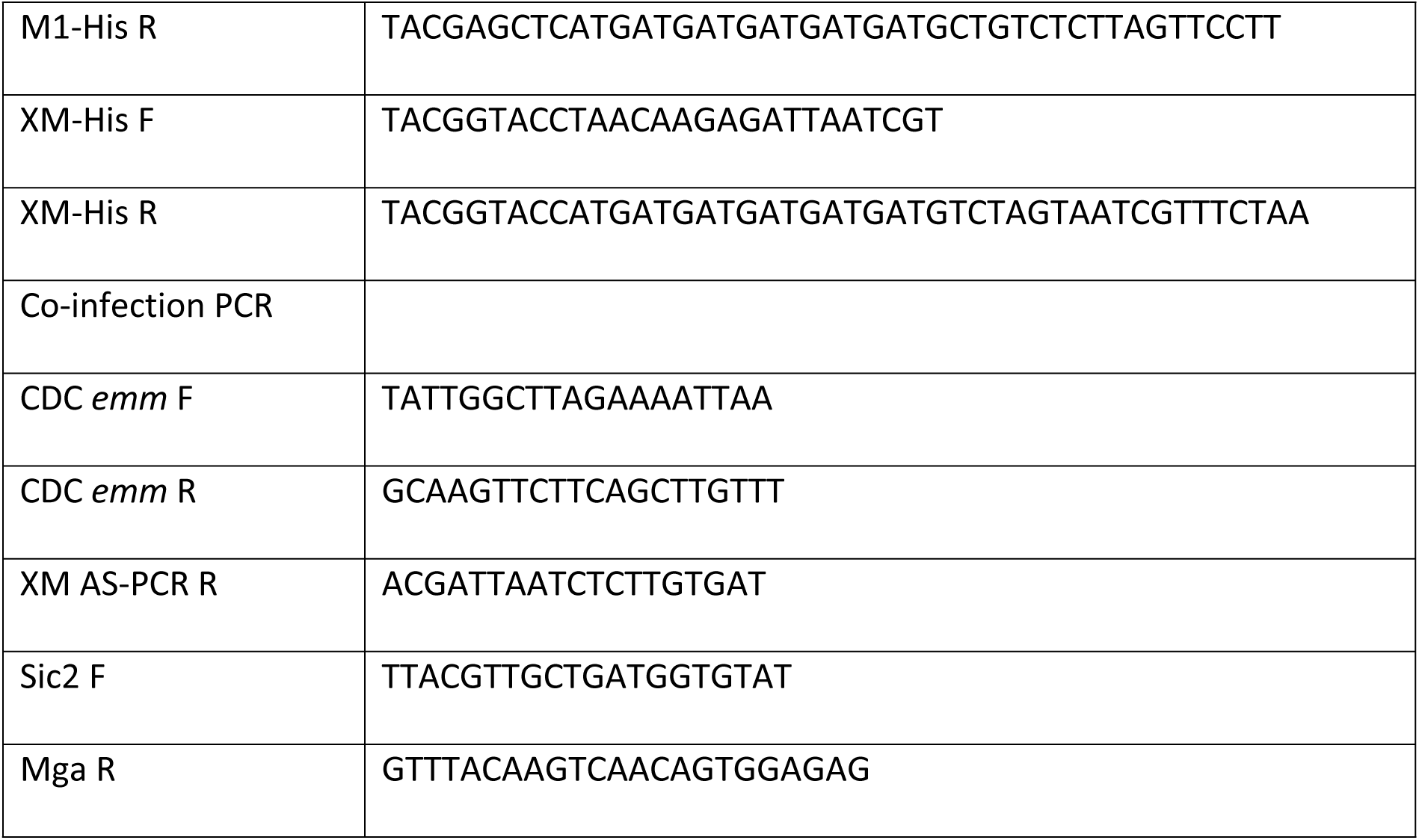
Primers used in current study.

### M1 protein detection by Western blot

Cell wall extracts and growth supernatant from *S. pyogenes* strains were prepared as previously described [46] prior to comparison by western blotting. Samples were separated by SDS-PAGE using 10% Bis-Tris gels (ThermoFisher Scientific) and transferred onto nitrocellulose. M1 protein was detected using polyclonal antiserum, diluted 1:1000, from mice or rabbits vaccinated with M1 HVR recombinantly produced in *E. coli* [46]. Secondary antibodies used were HRP-conjugated goat anti-mouse IgG and anti-rabbit IgG (Abcam), used at 1:80,000.

### Murine infection

Female FVB/n or BALB/c at 5-6 weeks were used for infection. For intramuscular infections, *S. pyogenes* grown on CBA plates for 16 h was washed and suspended in sterile PBS to give 2×10^8^ CFU per 50 μl dose. For co-infections, bacterial strains were mixed 1:1. A dose of 2×10^8^ CFU per 50 μl was also used for infections using *L. lactis*. Bacterial suspension was inoculated in the right hind limb and infection allowed to proceed for 3 or 24 h. Blood was collected by cardiac puncture and cultured for bacterial quantification. Tissues of interest were harvested and homogenised in sterile PBS to allow culturing of bacteria for quantification.

PCR was used to differentiate bacterial colonies following co-infection. H584_XM_ was detected using an allele-specific primer (AS-PCR). Up to ten colonies per organ were randomly selected, emulsified in 5% Chelex (Sigma Aldrich) suspended in molecular grade water and boiled for 10 minutes prior to PCR.

Intranasal infections were performed using 5-6 week old female FVB/n mice as previously described [22]. For upper respiratory tract infection, mice were infected with 5 × 10^6^ CFU in 5 µL and carriage assessed by nose press onto selective media over 96 h. Airborne shedding of *S. pyogenes* was observed by leaving streptococcal selective COBA (Colistin sulphate Oxolinic acid Blood Agar) plates face up in the top of each cage for a period of 4 hours at the same time each day during infection. Loss of colonisation was considered as two consecutive days with negative nose press plates for an individual mouse. At the end of infection, mice were culled, and blood collected by cardiac puncture, and selected tissues harvested and homogenised for bacterial culture and quantification. For lower respiratory tract infection mice were infected with 2.5×10^8^ CFU in 10 µL and bacterial burden in nasopharynx, liver and spleen enumerated after 72 h.

### Whole blood survival assays

Survival of *S. pyogenes* in whole human blood from healthy volunteers was assessed as previously described [46] with minor modifications. Heparinised blood was inoculated with 300-500 CFU of early logarithmic stage *S. pyogenes* (OD_600_ 0.15-0.17) prior to 3 h incubation at 37°C with end over end rotation. Bacteria within the blood was then cultured for quantification and multiplication factor determined as the number of CFU at 3 h divided by the number at 0 h. The same method was used to assess survival of *S. pyogenes* in blood from 6-week-old female BALB/c mice. Blood was collected by cardiac puncture into tubes containing heparin immediately prior to inoculation with *S. pyogenes*.

### Purification of recombinantly expressed M1 and XM protein in *L. lactis*

Recombinant M1 and XM protein were produced in *L. lactis*. The sequence for *emm1* was introduced into vector pOri23 [47] allowing expression on the cell wall of *L. lactis*. This vector was further modified by site directed mutagenesis using the QuikChange II kit (Agilent) to create two new vectors that would allow for secretion of His-tagged proteins, one containing a His-tag and stop codon directly upstream of the sequence for the LPXTG cell wall anchor and another containing a His-tag and stop codon in the spacer region sequence to emulate the truncation found in H584_XM_. The vectors were transformed into *L. lactis* MG1363 as previously described [48].

*L. lactis* bearing the vectors were cultured overnight as standard before collecting the growth supernatant and sterilising with 0.2 µm vacuum filter units (Millipore). Sterile supernatant was concentrated using 10 kDa MWCO centrifugal filter units (Amicon). His-tagged M1 and XM protein were purified by metal affinity chromatography using Ni-NTA resin and analysed by SDS-PAGE. Fractions containing recombinant protein were pooled and dialysed overnight at 4°C into PBS using Slide-A-lyser dialysis cassettes (ThermoFisher Scientific). Protein was quantified using Bradford assay or Qubit 4 fluorometer (ThermoFisher Scientific).

### Binding assays

The ability of *S. pyogenes* strains to adhere to various host factors was assessed using a solid-phase adherence assay, modified from [49]. The wells of a sterile 96-well flat bottom tissue culture plate were coated with 2.5 µg of human fibrinogen, fibronectin, collagen type I or collagen type IV (Sigma), prepared in carbonate-bicarbonate buffer, and incubated overnight at 4°C. Wells were washed three times with sterile PBS and then blocked with 100 µl 2% w/v bovine serum albumin (BSA) (Sigma) in PBS for 30 min at room temperature. Wells were washed again, and 100 µl of overnight cultures of *S. pyogenes* prepared to OD_600_ 10 were added to each well. Plates were incubated at 37°C for 2 h before staining with 50 µl 0.5% w/v crystal violet in distilled water. Wells were washed five times with PBS and incubated for 10 minutes with 5% v/v acetic acid before measuring OD_590_.

To assess the binding of recombinantly produced M1 protein and XM protein, 96-well high binding plates (Corning) were prepared by coating with 2.5 µg of human fibrinogen, fibronectin, collagen type I or collagen type IV as above. Triplicate wells were also coated with 100 μL recombinant M1 or XM protein, prepared to a concentration of 1.85 nM in carbonate-bicarbonate buffer. Plates were incubated at 4°C overnight, and the wells washed three times with PBS-Tween (0.05%). Wells were blocked for 1 hour at room temperature using 5% w/v skim milk powder, 0.1% Tween, 0.1% normal goat serum, in PBS. Washing was repeated three times before adding 100 μL recombinant M1 or XM (1.85 nM) to the wells coated with host factors and incubating for 2 h at room temperature. Wells were washed three times before 100 μL of anti-M1 HVR mouse serum (1:1000 dilution in blocking buffer) was added and wells incubated for 1 h at room temperature. Wells were again washed three times and 100 μL goat anti-mouse HRP conjugated antibody (diluted 1:80,000 in blocking buffer) was added and the plate incubated for 1 h at room temperature. The wells were washed three times before the addition of 50 μL TMB. The reaction was stopped with 50 μL H_2_SO_4_, and then absorbance read at 450 nm. Binding of the recombinant protein to the different substrates was reported as a binding ratio (A_450_ protein bound to substrate/A_450_ protein).

### Preparation and analysis of Biotinylated Glycosaminoglycan polysaccharide probes

Biotinylated HA-S350 from *Streptococcus pyogenes,* 350 kDa (LifeCore), CSA (from bovine trachea, Sigma), CSB (from porcine intestinal mucosa, Sigma), CSC (from shark cartilage, Sigma) and heparin (from porcine intestinal mucosa, Sigma) were prepared and purified essentially as described [50]. The level of biotin introduced was one for every 50 carboxyl groups [51]. In brief, 50 mg of GAG polysaccharide were dissolved in 0.1 M MES buffer (pH 5.5) before addition of 109 ml of 25 mM biotin LC-hydrazide (Pierce, Tattenhall, UK) solution in Me_2_SO and 130 ml of 25 mg/ml 1-ethyl-2-(2-dimethylamino propyl) carbodiimide solution in 0.1 M MES buffer. The reaction mixtures were stirred at room temperature for 24 h and dialyzed extensively against deionized water. A short Sephadex G-10 column (1.6 x 30 cm) was used to remove remaining biotin and other reagents. Quantitation of GAG polysaccharides and their modified forms was carried out by carbazole assay [52].

### Glycan microarray analyses

All glycan binding analysis was performed at the ICL Carbohydrate Microarray Facility. The generation of broad-spectrum NGL-based screening glycan arrays, the focused GAG oligosaccharide arrays and GAG polysaccharide arrays was performed essentially as previously described [53,54]. Details of the microarray preparation, methods used for binding assays and data analysis are provided in S2 Table, in accordance with MIRAGE (Minimum Information Required for A Glycomics Experiment) guidelines for reporting of glycan microarray-based data [55]. The procedures for microarray binding experiments with recombinant M proteins and whole bacterial cells are given below.

The analyses of His-tagged recombinant M1 protein (rM1) and truncated M1 protein (rXM) using the broad-spectrum NGL-based screening glycan arrays and the focused GAG oligosaccharide microarray were performed essentially as previously described [56] at ambient temperature. In brief, the His-tagged rM1 and rXM were diluted in 1 %(w/v) BSA in in HEPES buffered saline (HBS: 5 mM HEPES, pH 7.4, 150 mM NaCl) supplemented with 5 mM CaCl_2_ at a final concentration of 200 µg/mL and overlaid onto the arrays for a 1.5-hour incubation. The arrays were then incubated for 1 hour with a mixture of mouse anti-His antibody (Sigma SAB4200620), precomplexed at a ratio of 1:1 (by weight) with biotinylated anti-mouse IgG (Sigma B7264) and overlaid at 10 µg/mL. the final detection was with 30min overlay of streptavidin-Alexa Fluor 647 (Molecular Probes) at 1 µg/mL. After washing the slides were dried in a microarray slide spinner centrifuge before scanning and quantitation.

Microarray analyses of fluorescently labelled bacteria was performed as previously described [57]. Briefly, fluorescently labelled bacterial cultures at OD 1 were resuspended in HBS buffer supplemented with 5 mM CaCl_2_ and 0.01% Tween, at pH 7.5) and were used for overlays on glycan microarrays. 100 µL of bacterial suspension was applied to the incubation chamber with the microarrays, incubated for 1 hour at room temperature under mild agitation on an oscillating platform and washed three times with binding buffer followed by two washes with HPLC grade water to remove salts from the array. Slides were dried under a mild nitrogen flow and scanned for quantitation as described below.

For imaging scanning, quantitation and data analysis please see MIRAGE document (S2 Table).

### Analysis of capsule synthesis and transcription of capsule related genes

Hyaluronan capsule was extracted from bacterial subcultured to OD 0.3 (mid -logarithmic stage) in fresh THB or grown overnight on CBA. Samples were taken for CFU enumeration. A volume of 1.5 mL of culture was pelleted at 16,000 x g for 5 minutes and the resulting pellet resuspended in 1 mL of pH 7.0 Tris buffer, before centrifugation was repeated. The final bacterial pellet was resuspending in 400 µL buffer. To this bacterial suspension was added 0.72 U proteinase K, and samples incubated at 37°C for 15 minutes, before adding 1 mL phenol:chloroform:isoamyl alcohol (25:24:1 v/v/v) and mixing thoroughly. Samples were centrifuged for 10 minutes at 16,000 x g and the aqueous layer transferred to a new microcentrifuge tube and 1 mL of chloroform added and mixed well. Samples were centrifuged at 16,000 x g for 10 minutes and the aqueous layer transferred to a new tube and stored at −20°C until required. HA was quantified using the Hyaluronan DuoSet ELISA (R&D Systems) according to the manufacturer protocol. HA amount was normalised to CFU number.

Transcriptional analysis was carried out on *S. pyogenes* grown in THB for 24 h or from infected murine thighs tissue. Mice were infected intramuscularly for 24 h and thighs dissected and chopped. A 20 µl sample was taken and serially diluted for CFU enumeration. The tissue was pressed through a 0.7 µm cell sieve into a 50 ml tube on ice, the sieve was flushed with 10 ml ice cold PBS before tubes were spun at 2,500 x g for 10 min. Excess PBS was removed before proceeding directly to RNA extraction using hot phenol [58]. RNA pellets were air dried before resuspending in 50 µl molecular grade H_2_O and heated at 55°C to ensure all RNA was dissolved. Samples were treated with the Turbo DNA-free kit (Invitrogen). DNase treated RNA was then stored at −80°C. RNA samples were converted to cDNA using Superscript IV reverse transcriptase (Invitrogen). Quantitative PCR was carried out for the genes gyrA (F: 5’AGCGAGACAGATGTCATTGCTCAG, R: 5’CCAGTCAAACGACGAAACG), covS (F: 5’CGCTCAGATATTTCGTCAGG, R: 5’CGTAAATTCATGGCTGACATCAC), covR (F: 5’AAATATGATGAAGCCGTTGAGAC, R: 5’GCCACGCACTGTTTGGATAT), and hasA (F: 5’CCACATGACTATAAAGTTGCTG, R: 5’CTGATAACGGATAGGTCTGTG) using SYBR Green Jumpstart Taq ready mix (Sigma) [58]. A standard curve was generated using a plasmid containing a single copy of each target gene as previously described [58]. Reactions were thermocycled by the AriaMx qPCR cycler (Agilent) using the following cycle:

1. 95°C 3 min
2. 95°C 10 sec
3. 60°C 10 sec
4. 72°C 10 sec (Fluorescence measured)
5. Steps 2-4 repeated 39 times

The level of gene expression was determined for each gene using the AriaMx software (Agilent) and normalised by calculating transcript copy number per 1,000 copies of gyrA.

### Statistics

All statistical analyses were performed using GraphPad Prism 8.0. A p value of ≤ 0.05 was considered statistically significant. Sample size calculations were performed using GPower 3.1. Where appropriate, data from *in vivo* infections were normalised by Log_10_ transformations, and comparisons between two groups performed using t test, and comparisons between more than two groups performed using two-way ANOVA with Tukey’s test for multiple comparisons.

## Acknowledgments and Funding

SS acknowledges the support of the NIHR Biomedical Research Centre (BRC) awarded to Imperial College London (ICL). YL acknowledges John Hogwood, National Institute for Biological Standards and Control (NIBSC) for the heparin polysaccharides used in GAG microarray preparation. The authors would like to thank Mahrokh Nohadani and Professor Gordon Stamp for their expertise in histopathological preparation and analysis of specimens.

This work was funded by UKRI (Medical Research Council Training grant MR/R502376/1 and project grant MR/L008610/1) and in part by Wellcome Trust grant 215539/Z/19/Z. The glycan microarray studies were performed in the Carbohydrate Microarray Facility within the ICL Glycosciences Laboratory supported by the Wellcome Trust Biomedical Resource Grants (WT099197/Z/12/Z, 108430/Z/15/Z and 218304/Z/19/Z) and March of Dimes Prematurity Research Centre grant (22-FY18-82).

## Supporting Information

**S1 Fig. Growth of *S. pyogenes* and *L. lactis* strains were not affected by differences in expression of M1 protein.** *S. pyogenes* was sub-cultured and grown in THB in 5% CO_2_ and growth was checked hourly for 1-6 hours and again at 24 hours post sub-culture. No difference was seen in growth (A, n=3 biological replicates, mean and standard deviation shown). Growth of L. lactis strains over-expressing M1 or XM protein were compared to L. lactis containing just the empty pOri23 plasmid. Strains were grown in GM17 media supplemented with erythromycin and samples taken for enumeration hourly for 1-6 hours and again at 24 hours post sub-culture. No difference was seen (B, n=3 biological replicates, mean and standard deviation shown).

**S2 Fig. Mice were infected intramuscularly with *L. lactis* expressing cell wall anchored M1 protein (pOri23_emm1_), extracellular XM protein (pOri23_XM_), or simply harbouring the pOri23 plasmid with no additional gene (pOri23).** Expression of XM led to significantly decreased recovery of pOri23_XM_ from the thigh (CFU/100 mg) compared to pOri23_emm1_, but no difference in systemic (CFU/ml of blood, CFU/100 mg liver, CFU/whole spleen) or lymphatic dissemination (CFU/lymph node) was seen 3 h (A, n=6) or 24 h after infection (B, n=18, three pooled experiments) (** p<0.005, *** p<0.0005).

**S3 Fig. Broad-spectrum glycan microarray screening analyses of recombinant M1 and XM proteins.** (A) Histogram charts showing binding results of His-tagged M1 and XM. Binding was detected using a mouse anti-His antibody in conjunction with biotinylated anti-mouse IgG, followed by streptavidin-Alexa Fluor 647. Fluorescence intensities represent the average of duplicate spots (5 fmol per spot level), with error bars indicating half the difference between replicates. The glycan probes were arranged by glycan class in color-coded panels. (B) A detailed comparison of binding for each protein to heparin-related probes in the arrays. Full microarray results with probe list and probe sequences and complete binding data are provided in S1 File.

**S4 Fig. Glycosaminoglycan polysaccharide binding by *S. pyogenes* isogenic strains H584, H584_XM_ and H584_Δemm1_.** Heatmap showing the relative binding intensities to five biotinylated GAG polysaccharides of fluorescently labelled *S. pyogenes isogenic strains H584, H584_Demm1_ and H584_XM_.* The background subtracted mean fluorescence intensities of triplicate spots on the microarray is shown and the resulting values coloured as follows: Dark blue (0-1 %); Light blue (1-10 %); Yellow (10-30 %); Orange (30-70%); Red (70-100%); Grey, signals with high SD due to artifacts on the slide. *100%, the maximum binding score observed for a given strain*.

**S1 Table. Summary of full report by histopathologist**.

**S2 Table. Supplementary glycan microarray document based on MIRAGE Glycan Microarray guidelines (doi:10.3762/mirage.3).**

**S1 File. Appendix glycan array data.**

